# Characterization of Clinical *Fusobacterium nucleatum* Isolates from Oral Squamous Cell Carcinoma Patients

**DOI:** 10.1101/2025.01.08.631950

**Authors:** Serene Lim, Wan-Hsin Hsueh, Ni-Hung Wu, Claire Hodges, Christopher Vuong, Kynzi Smith, Jenn-Ren Hsiao, Jeffrey S Chang, Jang-Yang Chang, Jenn-Wei Chen, I-Hsiu Huang

**Author notes:** Correspondence: I-Hsiu Huang.

## Abstract

*Fusobacterium nucleatum* (*Fn*), a Gram-negative anaerobe primarily residing in the oral cavity, has garnered increasing attention for its role in a broad spectrum of human diseases. While typically absent or rarely detected outside the oral cavity in healthy individuals, *Fn* is frequently found at extra-oral sites under disease conditions and has been implicated in cancer progression and prognosis^1^. In oral squamous cell carcinoma (OSCC), the abundance of *Fn* significantly increases as the disease progresses, promoting cell invasion and metastasis. Furthermore, substantial evidence links *Fn* to accelerated tumor growth and metastatic progression in colorectal cancer (CRC), where its presence is also associated with chemotherapy resistance and poor prognosis. Here, to further elucidate the pathogenic mechanisms of *Fn* in cancer progression, this study characterized the physiological traits, virulence factor expression, and impacts on cancer cells of 10 *Fn* strains, including two well-characterized ATCC strains and eight clinical isolates. The clinical isolates consisted of three strains from saliva samples of 117 OSCC patients and five strains from 160 non-cancer individuals. Results indicated that oral isolates, regardless of disease origin, all belong to *Fn* subspecies *polymorphum*. The ATCC strains (23726 and 25586) exhibited shorter cell lengths and faster growth rates compared to the clinical isolates. Both ATCC strains formed stable biofilms and expressed key virulence genes, including *aim1*, *fadA*, *fomA*, and *radD*. All isolates tested showed sensitivity to a panel of eight different antibiotics. Interestingly, only one clinical isolate displayed similar stimulation of CRC cell migration as the two ATCC strains, while the other seven displayed no such ability. Collectively, these findings suggest that the most virulent strains, in terms of biofilm formation and virulence gene expression, are not necessarily the most pathogenic in the context of cancer cell interactions. Future pan-genomic analyses, incorporating whole-genome sequencing of the clinical isolates, will aim to delineate the genetic determinants contributing to the carcinogenic potential of *Fn*.

## Introduction

Recent advances in microbial detection have highlighted the role of previously overlooked microorganisms in human diseases. One such emerging pathogen is *Fusobacterium nucleatum* (*Fn*), a spindle-shaped, anaerobic bacterium originally regarded as a harmless oral commensal. However, its presence in inflammatory conditions and systemic diseases, particularly its ability to thrive in tumor microenvironments, underscores its role in cancer progression^1–3^. Phylogenetic analyses identify four *Fn* subspecies—*animalis*, *nucleatum*, *polymorphum*, and *vincentii*—each with distinct disease associations^4–6^. For example, *polymorphum* and *vincentii* are common in healthy oral cavities, while *nucleatum* is linked to periodontal disease and systemic infections^6^. *Animalis*, predominantly found in the colon, contributes to colorectal cancer (CRC) development and intestinal inflammation, while *vincentii* is associated with prostate inflammation and cancer^7,8^. These variations reflect the genetic diversity and pathogenic potential of *Fn* subspecies.

Once seen as benign, *Fn* is now recognized as a contributor to numerous diseases, including periodontal disease, atherosclerosis, diabetes, rheumatoid arthritis, adverse pregnancy outcomes, and Alzheimer’s^2,9–15^. Notably, its association with cancers such as oral squamous cell carcinoma (OSCC), CRC, and malignancies of the stomach, esophagus, pancreas, and breast highlights its significant oncogenic potential. Studies reveal that *Fn* abundance increases with OSCC progression, with *polymorphum* identified as the predominant subspecies in biopsy samples^16^. *Fn* contributes to OSCC through oncogenic signaling pathways, including activation of β-catenin and Wnt5a, which drive cell proliferation, invasion, and resistance to treatments such as cisplatin^17–19^. It also facilitates cancer progression by promoting lactate production and immune evasion^20,21^.

CRC, the second leading cause of cancer-related mortality, is similarly influenced by *Fn*. Frequently detected in CRC tissues, its levels correlate with the progression from benign adenomas to carcinomas^22–24^. Key virulence factors such as *FadA* and *Fap2* play critical roles in this process. *FadA* binds to E-cadherin on colonic epithelial cells, disrupting cell junctions and activating oncogenic pathways, while *Fap2* inhibits natural killer (*NK*) cell activity, enabling immune evasion^17,25–27^. *Fn* also enhances biofilm formation and induces chemoresistance, particularly against 5-fluorouracil, further contributing to CRC progression and poor patient outcomes^28–30^.

Despite its established role in cancer, significant genetic and phenotypic variability among *Fn* strains complicates research. Only certain strains possess the specific carcinogenic traits necessary for tumor promotion, and with over 2,000 genes in its genome, many virulence factors remain unidentified^31,32^. Understanding *Fn*’s mechanisms in cancer is crucial for developing targeted therapies. In the present study, we aimed to address these gaps by characterizing the phenotypic and genotypic differences among *Fn* strains from OSCC patients and exploring novel virulence factors to inform future therapeutic strategies.

## Materials and Methods

### Bacterial strains and cultures

*F. nucleatum* strains 23726 and 25585 were purchased from American Type Culture Collection (ATCC, Manassas, Virginia). Clinical strains were isolated from National Cheng Kung University Hospital (NCKUH) patients’ saliva (IRB number: B-ER-106-234). Saliva samples were diluted 1:100 and 1:1000 in phosphate-buffered saline and plated onto Crystal Violet-Erythromycin (CVE) agar plates. The plates were incubated in an anaerobic workstation (Coy Laboratory Products, MI) anaerobically (5% CO2, 5% H2, 90% N2) at 37°C for at least 72 hours. Colonies with rounded edges and a purple-grey hue were selected for further analysis. These colonies were subjected to Gram staining for morphological screening, where spindle-shaped, Gram-negative bacteria were identified. The selected colonies were then streaked onto fresh CVE agar plates for further purification. Species confirmation was performed via PCR amplification of the Fn 16S rRNA gene with primers Fn16s-F (5’-GAGAGAGCTTTGCGTCC-3’) and Fn16s-R (5’-TGGGCGCTGAGGTTCGAC-3’). The PCR program was set as follows: 95℃ for 5 min, 35 cycles of 95 ℃ for 30 s, 55 ℃ for 30 s, and 72 ℃ for 30s and a final extension at 72℃ for 5 min. Following species and sub-species confirmation by Sanger sequencing, bacterial pellets were harvested by centrifugation at 16,000 g for 1 minute, resuspended in LaboBanker 1 (Kurabo, Japan), and stored at −80°C until further analysis. TSPC (Tryptic soy Broth (TSB) (BD, USA) supplemented with 0.1% (w/v) L-cysteine (Thermo Fisher Scientific, USA), 1% (w/v) bacto peptone (BD, USA) was used for *F. nucleatum* culturing at 37°C anaerobically.

### Cell Morphology and growth curve analysis

Fn strains were retrieved from −80°C storage and streaked onto TSPC agar. The plates were then incubated at 37°C anaerobically for 48 hours. After incubation, bacterial colonies were harvested and smeared onto glass slides for Gram staining and subsequent imaging. Images were captured by confocal microscopy (Leica, Germany) from at least three distinct regions per sample, and measurements of three individual cells per region were taken to ensure robust data collection. Cell length measurements were conducted under 1,000x oil immersion magnification using LAS X software (Leica, Germany). Imaris software (Oxford Instruments, UK) was used to analyze the volumetric dimensions of individual bacterial cells, with false fluorescence signals filtered out to ensure data accuracy.

A 48-hour-old bacterial colony was inoculated into TSPC broth and cultured anaerobically at 37°C overnight. After incubation, 1 ml of the culture was transferred to 9 ml of fresh TSPC medium and incubated for 17 hours under the same conditions. The optical density at 600 nm (OD600) of each refreshed culture was then adjusted to 0.01 using a Novaspec III+ spectrophotometer (Biochrom, UK). The bacterial suspension was divided into three glass test tubes, each containing 10 ml of culture with an OD600 of 0.01. These test tubes were incubated anaerobically at 37°C, and OD600 readings were taken at 1-hour intervals using an Ultraspec10 spectrophotometer (Biochrom, UK) until the stationary phase was reached, which occurred after approximately 16 hours. Fresh TSPC broth was used as a blank for spectrophotometric measurements. This process was repeated in three independent experimental trials to ensure reliability and reproducibility of the results.

### Biofilm formation assay

Fn strains were retrieved from −80°C storage and streaked onto TSPC agar. The plates were then incubated at 37°C anaerobically for 48 hours. After incubation, single colonies were transferred to 10 ml of TSPC medium and grown anaerobically at 37°C for 17 hours. To initiate biofilm formation, the OD600 of each overnight culture was adjusted to 0.1 in fresh TSPC broth. A 1 ml aliquot of this bacterial suspension was added to each well of 24-well polystyrene plates (Falcon, USA) and incubated statically for 96 hours at 37°C under anaerobic conditions to allow biofilm formation. Following the 96-hour incubation, biofilm quantification was performed using the crystal violet (CV) staining method. Non-adherent cells were removed by aspirating the spent medium, and 400 μl of 0.01% (w/v) CV solution (Alfa Aesar, USA) was added to each well to stain the biofilms. After 15 minutes at room temperature, the CV solution was discarded, and each well was carefully washed twice with 400 μl of sterile water to remove excess dye. To solubilize the stain, 500 μl of 30% (v/v) glacial acetic acid (J.T. Baker, USA) was added to each well, followed by gentle shaking for 15 minutes. The biofilm mass was then measured by reading the OD592 using a BioTek Synergy 2 microplate reader (Agilent, USA; Gen5 software v3.09). Wells containing only TSPC broth underwent the same staining process and served as baseline controls for calculating relative biofilm formation. Each experiment was performed in quintuplicate to ensure reproducibility.

### Antimicrobial susceptibility testing (AST)

Two-day-old bacterial colonies were transferred into 10 ml of TSPC medium and cultured anaerobically at 37°C for 17 hours. The overnight cultures were standardized to the 1 McFarland turbidity standard, and 1 ml of the bacterial suspension was evenly spread on TSPC agar plates. After 15 minutes of air drying in a biosafety cabinet, a single E-strip (Liofilchem, Italy) was applied to the center of the agar surface using sterilized forceps. The E-test strips, stored at −20°C, were allowed to reach room temperature for at least 30 minutes before use. Plates were incubated anaerobically at 37°C and read after 48-72 hours, depending on the clarity of the inhibition zones. The MIC was determined as the concentration at which the elliptical zone of inhibition intersected the E-strip, in accordance with Clinical and Laboratory Standards Institute (CLSI) guidelines.

### Virulence gene detection by PCR

Fn cultures were grown anaerobically for 17 hours and harvested by centrifugation at 13,000 g for 2 minutes. Genomic DNA was extracted using the Promega Wizard™ Genomic DNA Purification Kit (Promega, USA). Briefly, bacterial pellets were resuspended in 600 µl of Nuclei Lysis Solution, incubated at 80°C for 5 minutes, and treated with RNase A (Thermo Fisher Scientific, USA) at 37°C for 1 hour to remove RNA. Protein precipitation was initiated by adding 200 μl of Protein Precipitation Solution, followed by vortexing at high speed for 20 seconds. The samples were chilled on ice for 5 minutes and centrifuged at 13,000 g for 3 minutes to pellet the proteins. The supernatant containing genomic DNA was transferred to a clean tube, and DNA was precipitated by adding 600 μl of isopropanol. After centrifugation at 13,000 g for 2 minutes, the DNA pellet was washed with 600 μl of 70% ethanol, air-dried at room temperature for 10 minutes, and resuspended in 100 μl of sterile water. The DNA was solubilized at 65°C for 10 minutes and stored at −20°C until further use. Prior to PCR, the concentration and purity of the genomic DNA were measured using a NanoDrop OneC spectrophotometer (Thermo Fisher Scientific, USA), ensuring an A260/280 ratio of 1.8–2.0 and an A260/230 ratio of 2.0–2.2. DNA integrity was confirmed by electrophoresis (Thermo Fisher Scientific, USA) at 100V for 30 minutes on a 1% (w/v) agarose gel (Acros Organics, Belgium) prepared with 1x Tris-acetate-EDTA (TAE) buffer. Gels were stained with SYBR Safe DNA gel stain (Invitrogen, USA) and visualized under UV light using a Gel Doc XR+ imaging system (BioRad, USA) with Image Lab software (BioRad, USA). Gene-specific primers for each virulence factor were synthesized (Table 1). PCR reactions were performed in 50 µL volumes, consisting of 25 µl of Taq 2x master mix (New England Biolabs, USA), 2µl each of forward and reverse primers (10mM) (Integrated DNA Technologies, USA), 2µl of genomic DNA template, and 19µl of sterile water. Amplification was conducted using a SimpliAmp Thermal Cycler (Applied Biosystems, USA) with an initial denaturation at 95°C for 15 minutes, followed by 35 cycles of denaturation (95°C for 30 seconds), annealing (52–54°C for 90 seconds), and extension (72°C for 45 seconds). A final elongation step at 72°C for 7 minutes ensured complete amplification. Negative controls, lacking DNA template, were included in each batch to check for contamination. The amplified DNA fragments were separated by gel electrophoresis.

**Table 1.**
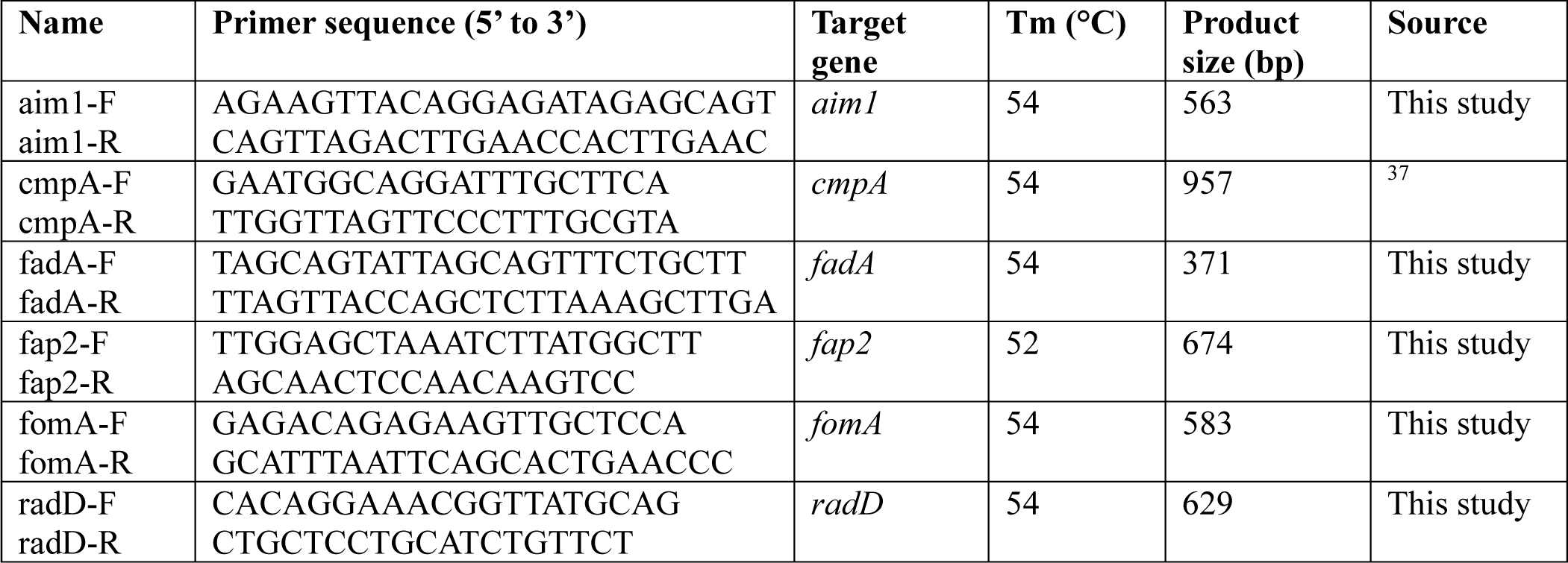
Primers used in this study.

### Cell lines and cultures

Human colon carcinoma cells HCT116 were purchased from ATCC (Manassas, VA). HCT116 cells were grown in McCoy’s 5A medium (Sigma-Aldrich, USA), with both media supplemented with 10% fetal bovine serum (MP Biomedicals, USA). The cells were kept in a humidified incubator at 37°C with 5% CO2. Routine maintenance involved gentle trypsinization using Trypsin-EDTA (Sigma-Aldrich Fine Chemicals Biosciences, USA), followed by reseeding. To prevent phenotypic drift, cells were not passaged more than 20 times.

### Cancer cell migration assay

To evaluate the impact of *Fn* isolates on cancer cell migration, a wound healing assay was performed using the CRC cell line. HCT116 cells were seeded at a density of 2×10^6^ cells per well in 6-well plates (Falcon, USA) and incubated overnight to achieve over 80% confluency. The next day, a wound gap was created by scratching the cell monolayer with a sterile P200 pipette tip (Axygen, USA). The medium and any detached cells were removed, and the remaining adherent cells were exposed to *Fn* isolates at an MOI of 10 under standard conditions. Wound closure was monitored by capturing images at 12 distinct regions per scratch using an inverted phase-contrast microscope (Leica, Germany). Images were taken immediately after the scratch (0 hours) and at subsequent time points (24 and 48 hours) at the same positions. The gap distance was measured by comparing wound closure between 0 and 48 hours. Each experiment was performed in triplicate to ensure statistical validity.

## Results

### *F. nucleatum* clinical isolates exhibits diversity in cell length

*Fusobacterium nucleatum* (*Fn*) isolates were obtained from the saliva samples of OSCC and non-OSCC patients at *NCKUH*. After PCR confirmation using *F. nucleatum*-specific primers, the isolates were submitted for Sanger sequencing to confirm their identity. For this preliminary study, we selected three OSCC and five non-OSCC isolates for all downstream analyses. Sanger sequencing results indicated that all eight clinical isolates shared the highest sequence homology to *F. nucleatum* subspecies *polymorphum* (supplementary data not shown). As numerous reports have indicated that the commonly used *F. nucleatum* strains 23726 and 25586 display shorter cell lengths compared to “wild-type” isolates, we subjected these isolates to cell morphology analysis. Microscopic imaging confirmed that all eight isolates were Gram-negative, fusiform-shaped rods. Cell length measurements revealed significant variation across strains, with sizes ranging from approximately 2 µm to 13 µm (Figure 1)—more than a fourfold difference. Among the isolates tested, isolate *Fn33399* exhibited the longest cell length (11–13 µm), while the ATCC strain 23726 had the shortest (2–4 µm). Notably, aside from isolates *Co31000* and *Co25544*, most *Fn* isolates tested measured over 7 µm, suggesting a potential size-related trend common to *Fn* isolates in saliva, regardless of clinical origin.

**Figure 1.**
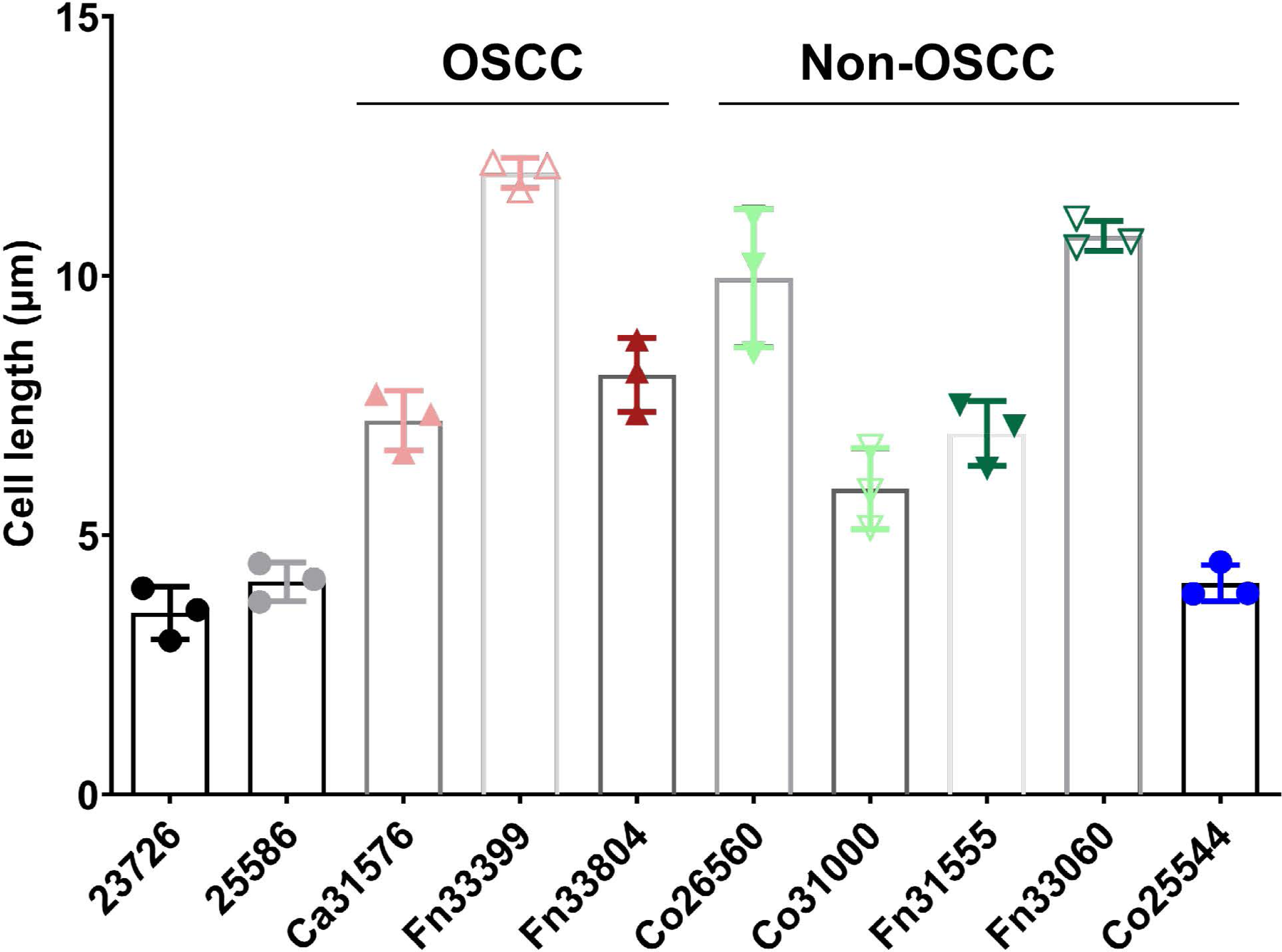
Cell length measurement of *F. nucleatum* isolates. Cell length of Fn ATCC strains (23726 and 25586) and eight clinical isolates were examined using brightfield microscope after gram staining. Cell length measurements were taken under 1,000x oil immersion magnification. Fn strain 23726 was used as the reference for statistical comparisons across all isolates. Statistical significance was evaluated using one-way ANOVA, with significance levels represented as follows: ns (not significant) p > 0.05, * p < 0.05, *** p < 0.001, **** p < 0.0001.

### *F. nucleatum* clinical isolates display diversity in growth rates

Distinct growth profiles, including lag, exponential, and stationary phases, were observed among the *Fn* isolates (Figure 2A). Compared to the wild-type strains 23726 and 25586, the growth rates of all other isolates were either similar or slower, with none exhibiting a faster rate. Based on the time required to reach the mid-exponential growth phase (*OD600* ∼1.0), the isolates were categorized into three groups. Fast-growing group (Figure 2B): Reaching mid-exponential phase within 11–14 hours, including *Fn* ATCC 23726, ATCC 25586, Fn33399, Fn33060, and *Co25544*. Slow to moderate-growing group (Figure 2C): Reaching mid-exponential phase within 14–20 hours, including isolates *Fn33804* and *Co31000*. Very slow-growing group (Figure 2D): Reaching mid-exponential growth within 20–26 hours, including isolates *Co31576*, *Co26560*, and *Fn31555*. Both OSCC and non-OSCC isolates demonstrated variability in growth kinetics, with no clear trend distinguishing the two groups.

**Figure 2.**
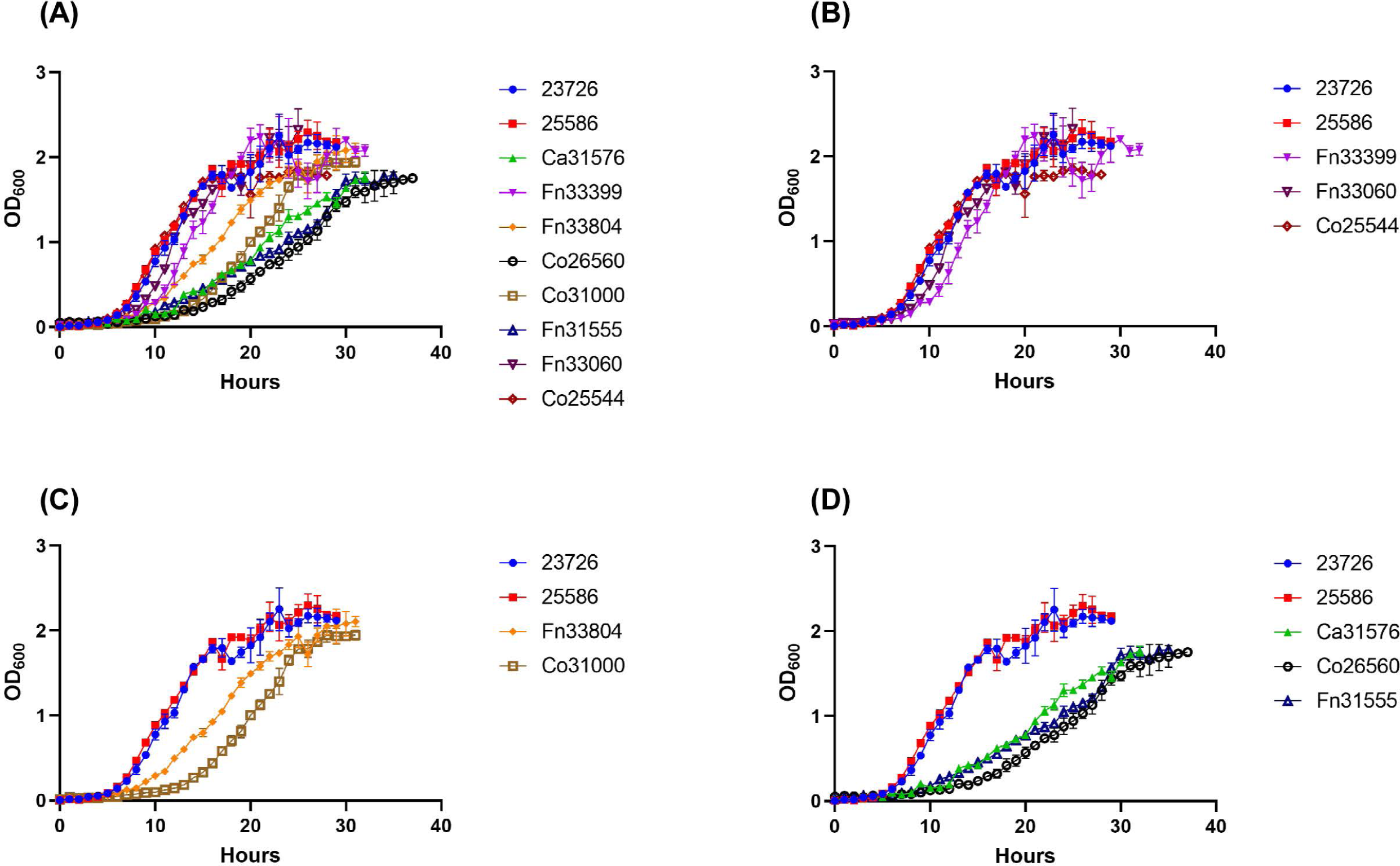
Planktonic growth kinetics of *F. nucleatum* isolates. Growth curves for Fn ATCC strains (23726 and 25586) and clinical isolates were monitored by measuring OD600 at 1-hour intervals until the stationary phase was achieved. All isolates were initially inoculated at an OD600 of 0.01. Based on the time required to reach the mid-exponential phase (OD600 ∼1.0), the isolates were classified into four distinct groups: (B) fast-growing, (C) slow to moderate growing, (D) very slow growing. Strain 23726 served as the reference for comparisons across all isolates. Data are presented as the mean ± SD from three independent experiments, each conducted in triplicate.

### Antimicrobial Susceptibility Testing

Eight antimicrobial agents were selected based on their mechanisms of action and chemical structure. These included macrolides (azithromycin), amphenicols (chloramphenicol), lincosamides (clindamycin), aminoglycosides (kanamycin), nitroimidazoles (metronidazole), fluoroquinolones (moxifloxacin), beta-lactams (penicillin G), and tetracycline. The minimum inhibitory concentration (*MIC*) results for the eight antibiotics tested against *Fn* isolates are presented in Table 2. All isolates, regardless of clinical background (ATCC strains, OSCC, non-OSCC), were susceptible to most antibiotics tested except for kanamycin. MIC values for azithromycin ranged from 0.75 to 6 µg/mL. Although *CLSI*-approved breakpoints for azithromycin in anaerobes are unavailable, it demonstrated the highest efficacy among macrolides against *Fn* (ref). MIC values for chloramphenicol ranged from 0.5 to 4 µg/mL, with the two ATCC strains (23726 and 25586) exhibiting higher resistance, though not yet reaching the breakpoint. MIC values for clindamycin ranged from 0.047 to 0.75 µg/mL, with clinical isolate Fn33804 at 0.75 µg/mL coming close to but not exceeding the *CLSI* breakpoint. Kanamycin exhibited the least effect, with MIC values ranging from 16 to 128 µg/mL. Both metronidazole and moxifloxacin exhibited strong inhibitory activity, with MIC values for metronidazole ranging from <0.016 to 0.094 µg/mL, far below the *CLSI* breakpoint of 8 µg/mL, and MIC values for moxifloxacin ranging from 0.047 to 0.25 µg/mL, well below the breakpoint of 2 µg/mL. Penicillin G showed strong inhibition, with MIC values ranging from 0.008 to 0.094 µg/mL, and MIC values for tetracycline ranged from 0.032 to 0.75 µg/mL. None of the tested isolates displayed resistance to the antibiotics tested.

**Table 2.** In <u>vitro</u> activity of antimicrobial agents against *F. nucleatum* isolates.

| ($\mu\text{g/ml}$) | AZM<sup>a</sup> | CHL<sup>b</sup><br>(CLSI<sup>i</sup> ≤<br>8) | CLI<sup>c</sup><br>(CLSI ≤<br>2) | KAN<sup>d</sup> | MTZ<sup>e</sup><br>(CLSI ≤<br>8) | MFX<sup>f</sup><br>(CLSI ≤<br>2) | PEN<sup>g</sup><br>(CLSI ≤<br>0.5) | TCY<sup>h</sup><br>(CLSI ≤<br>4) |
| --- | --- | --- | --- | --- | --- | --- | --- | --- |
| ATCC23726 | 4 | 4 | 0.125 | 128 | <0.016 | 0.094 | 0.032 | 0.064 |
| ATCC25586 | 3 | 2 | 0.064 | 48 | <0.016 | 0.25 | 0.047 | 0.047 |
| **<u>Oral cancer</u>** |  |  |  |  |  |  |  |  |
| Ca31576 | 2 | 0.75 | 0.47 | 32 | 0.032 | 0.125 | 0.094 | 0.38 |
| Fn33399 | 1 | 0.75 | 0.094 | 16 | 0.064 | 0.19 | 0.016 | 0.25 |
| Fn33804 | 1 | 1.5 | 0.75 | 96 | 0.094 | 0.125 | 0.008 | 0.19 |
| **<u>Non-cancer</u>** |  |  |  |  |  |  |  |  |
| Co26560 | 0.75 | 1 | 0.38 | 64 | 0.064 | 0.125 | 0.008 | 0.25 |
| Co31000 | 1.5 | 1 | 0.38 | 64 | 0.064 | 0.19 | 0.023 | 0.25 |
| Fn31444 | 0.75 | 1 | 0.064 | 64 | 0.023 | 0.19 | 0.016 | 0.19 |
| Fn33060 | 2 | 0.75 | 0.125 | 24 | 0.023 | 0.047 | 0.094 | 0.032 |
| Co25544 | 6 | 0.5 | 0.047 | 48 | <0.016 | 0.125 | 0.016 | 0.75 |

<sup>a</sup>AZM – Azithromycin. <sup>b</sup>CHL – Chloramphenicol. <sup>c</sup>CLI – Clindamycin. <sup>d</sup>KAN – Kanamycin. <sup>e</sup>MTZ – Metronidazole. <sup>f</sup>MFX – Moxifloxacin. <sup>g</sup>PEN – Penicillin G. <sup>h</sup>TCY – Tetracycline. <sup>i</sup>CLSI – Clinical and Laboratory Standards Institute.

### *F. nucleatum* clinical isolates appear to lack many known virulence genes

Six virulence genes were assessed in this study (Figure 3). Primers for each virulence gene were designed based on the genome sequence of *Fn* strain 23726 (Table 1). PCR results revealed that *aim1*, *cmpA*, *fap2*, and *radD* were detected only in the two control strains (23726 and 25586). None of the eight clinical isolates harbored these genes. Conversely, *fadA* was detected in all eight clinical isolates, and seven of eight clinical isolates harbored *fomA*, with the exception of isolate Co25544.

**Figure 3.**
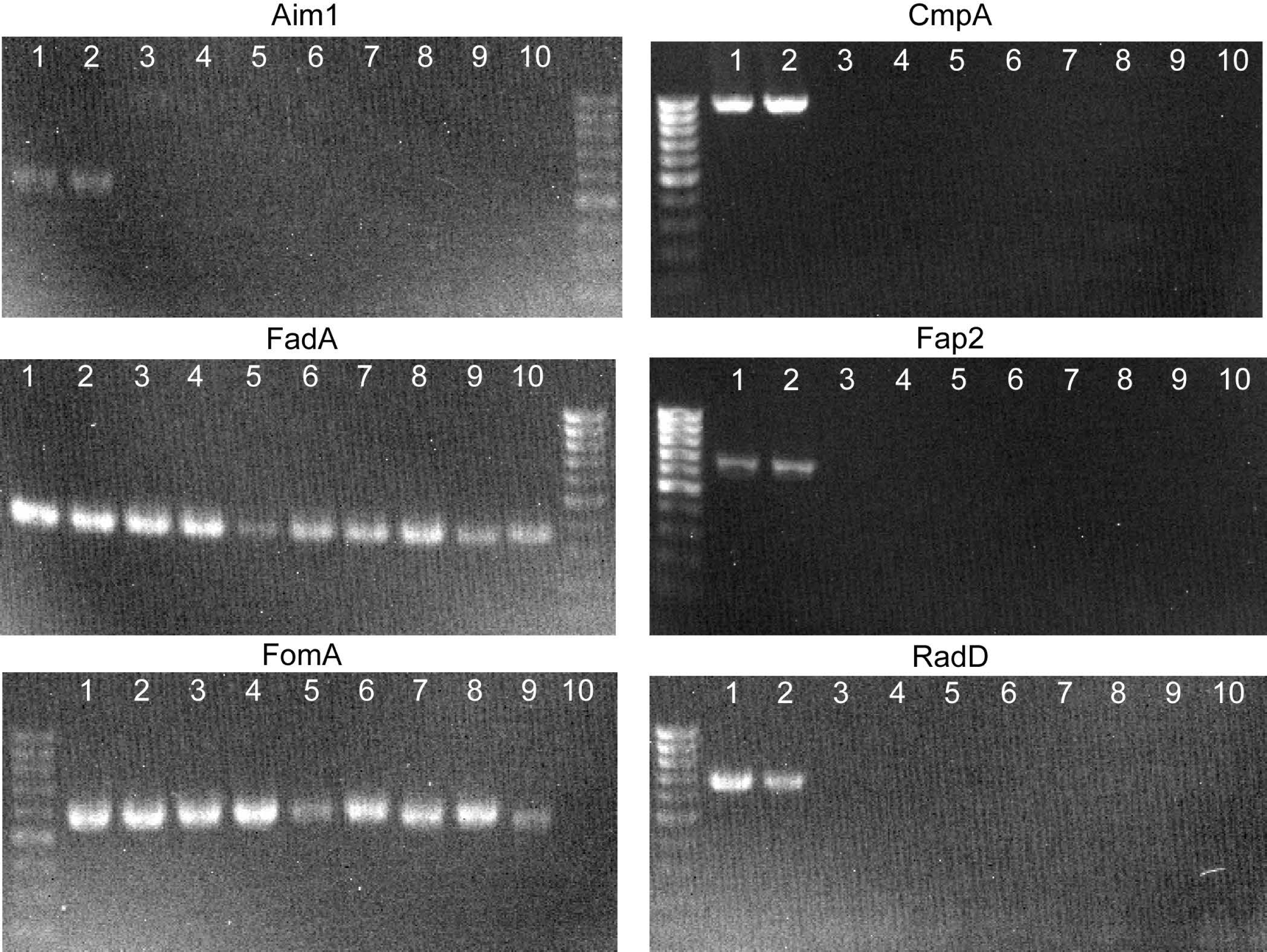
Detection of virulence genes in *F. nucleatum* isolates. PCR amplification results for the detection of aim1, *cmpA, fadA, fap2, fomA*, and *radD* in Fn ATCC strains (23726 and 25586) and clinical isolates. Genomic DNA was extracted from each strain and used as a template for amplification. Primers were designed based on the gene-specific sequences of the ATCC 23726 reference strain. 1-ATCC 23726, 2 - ATCC 25586, 3 - Ca31576, 4 – Fn33399, 5 – Fn33804, 6 – Co26560, 7 – Co31000, 8 – Fn31555, 9 – Fn33060, 10 – Co25544.

### Biofilm Formation Abilities

The biofilm-forming ability of *Fn* isolates was evaluated on abiotic surfaces using 96-well microplates and quantified via crystal violet (*CV*) staining. As shown in Figure 4, the two control strains (23726 and 25586) generally formed more robust biofilms than the eight clinical isolates, with strain 25586 exhibiting the highest biofilm-forming capacity. Most clinical isolates produced significantly less biofilm compared to the control strains. Comparisons between OSCC and non-OSCC groups showed no significant differences in biofilm production.

**Figure 4.**
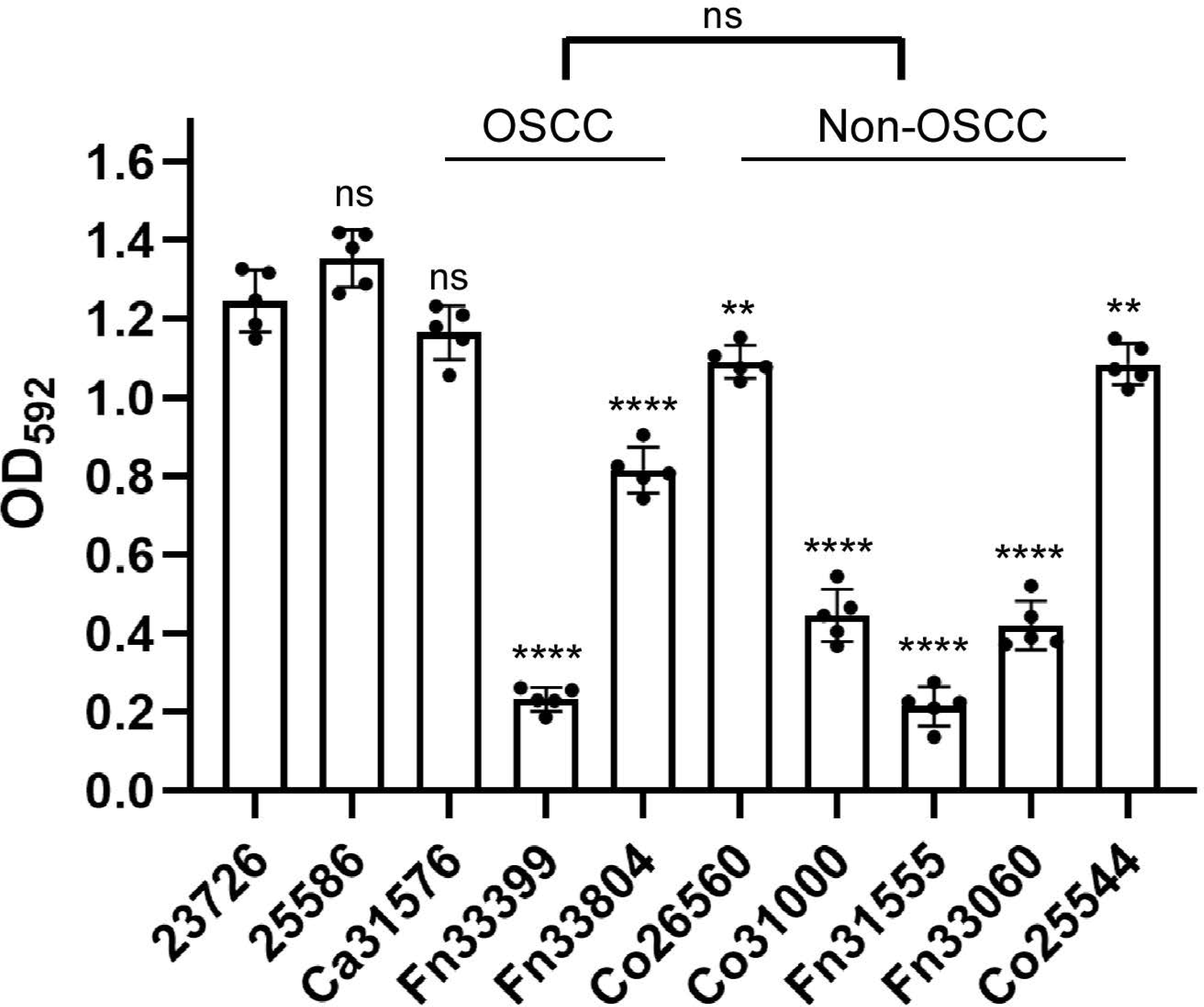
Biofilm formation abilities of *F. nucleatum* isolates. Biofilm formation was assessed for (A) Fn ATCC strains (23726 and 25586) and clinical isolates. All samples were inoculated at an OD600 of 0.1 and incubated statically for 96 hours. Biofilm quantification was performed using crystal violet (CV) staining, and the CV intensity was measured at 592 nm. Strain ATCC 23726 served as the reference for statistical comparisons across all isolates. Statistical significance was determined using one-way ANOVA, with results represented as follows: ** p < 0.01, **** p < 0.0001.

### Ability of *F. nucleatum* Isolates to Stimulate CRC **Cell Migration**

To evaluate the impact of *Fn* isolates on cancer cell migration, a wound healing assay was performed using the human CRC cell line *HCT116* (Figure 5). As expected, wound closure was significantly enhanced (*p* < 0.01) when *HCT116* cells were co-incubated with the ATCC strains 23726 and 25586. Strain 23726 was the most effective, promoting wound closure by 69%. Among the eight clinical isolates tested, only *Co25544* significantly enhanced cell migration, while the others did not exhibit this ability. These findings suggest that not all *Fn* isolates can stimulate CRC cell migration.

**Figure 5.**
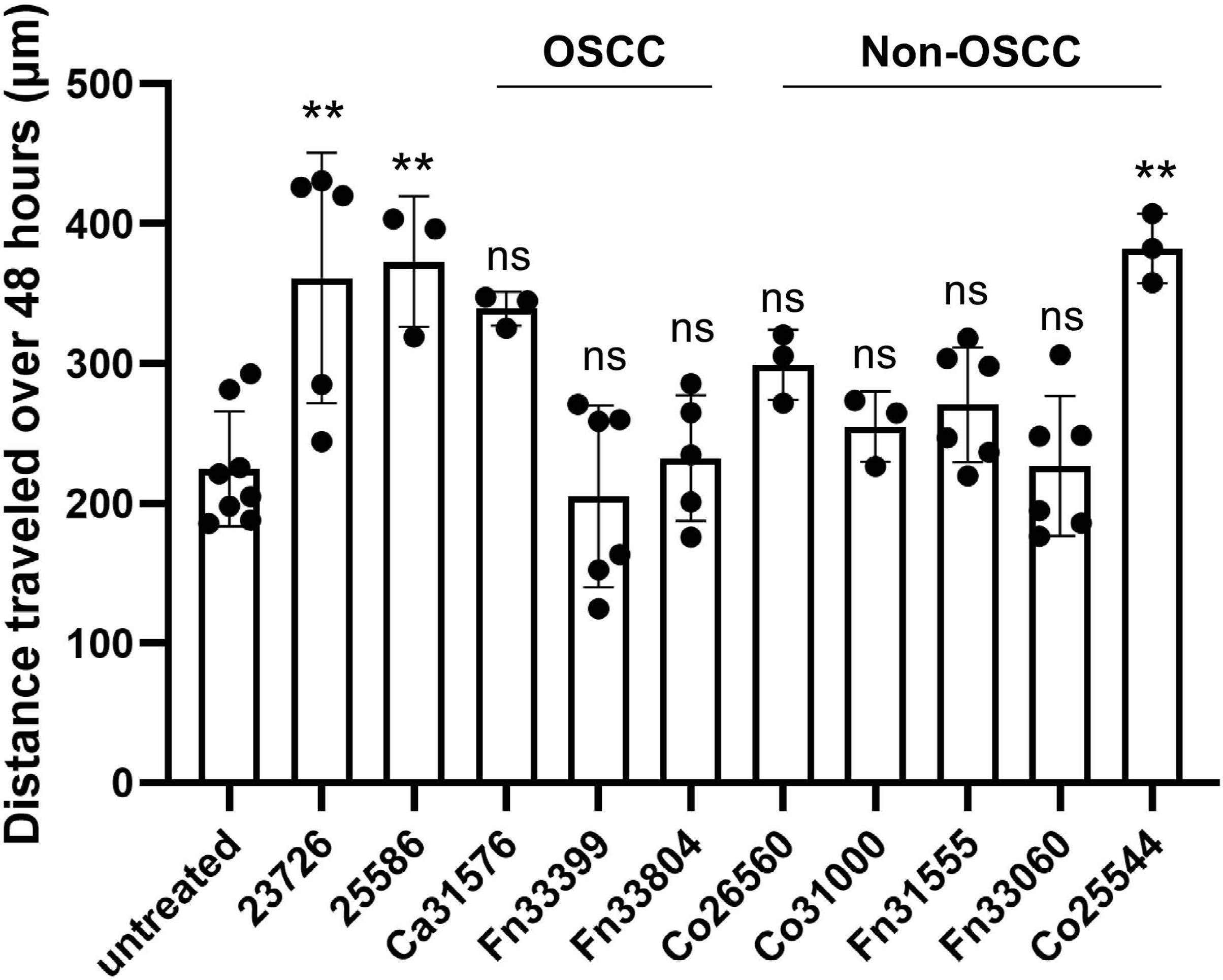
Motility of CRC cells co-incubated with *F. nucleatum* isolates. HCT116 cells were co-cultured with Fn ATCC strains (23726 and 25586) and clinical isolates at an MOI of 10 over a period of 48 hours. Gap closure was quantified by calculating the difference between the initial gap distance at 0 hours and the gap distance at 48 hours. Statistical significance was assessed using one-way ANOVA: ns p > 0.05, ** p < 0.01.

## Discussion

Human saliva is a valuable biofluid for clinical diagnostics and risk assessment due to its non-invasive and painless collection methods. Moreover, the oral environment generally maintains a relatively stable state over time, with only minor fluctuations caused by factors such as diet or infections. Importantly, the dominant oral microbiota in individuals is host-specific, further enhancing its diagnostic potential. Although multiple *F. nucleatum* isolates were obtained from the saliva of 277 enrolled patients, we included eight strains in this pilot study. Additionally, two well-characterized Fn ATCC strains (23726 and 25586) were included for comparative analysis.

Our results indicate that the majority of oral isolates, regardless of disease origin, belong to the *F. nucleatum* subspecie *polymorphum*, consistent with previous findings^16,33^. During invasive systemic infections, smaller bacterial cells tend to exhibit greater virulence^34^. In our study, the size of invasive Fn isolates ranged from 2 µm to 11 µm, with the ATCC strains 23726 (2–4 µm) and 25586 (3–5 µm) being on the smaller end, while the clinical isolates tended to measure longer. Anecdotal observations have been made in the past in which the two commonly used strains 23726 and 25586 are significantly shorter in length than “wild” isolates, perhaps due to repeated passage within laboratory setting. This suggests that shorter cell length may confer enhanced invasion capability, but longer cells are not necessarily incapable of invasion.

Bacterial morphology can also influence colonization of host niches^34,35^, with length potentially aiding biofilm formation. However, despite their shorter cell length, both ATCC strains displayed the greatest biofilm-forming capacities. Interestingly, our PCR analysis revealed that the clinical isolates lacked many genes associated with biofilm formation. For example, none of the isolates appeared to carry *cmpA* or *radD*. This finding suggests that robust biofilm formation might require other, as-yet-undiscovered factors. Additionally, these results challenge the assumption that known biofilm-associated genes are universally required across Fn strains.

While biofilms enhance bacterial resistance to antimicrobial agents, most Fn strains in this study, despite strong biofilm-forming capabilities, were susceptible to antibiotics, including chloramphenicol, clindamycin, metronidazole, moxifloxacin, and penicillin G. This susceptibility highlights the potential for effective therapeutic intervention even in biofilm-forming strains.

The ability of a pathogen to establish infection can also be influenced by bacterial growth rates, particularly for a strict anaerobe requiring specific atmospheric conditions. Both ATCC strains and clinical isolates Fn33399, Fn25749, Fn33060, and Fp Co25544 demonstrated the fastest growth, reaching mid-exponential phase within 11– 14 hours (OD600 ∼0.01). Other isolates required 16–26 hours to reach mid-exponential growth, potentially influencing their pathogenicity and colonization potential.

Virulence genes play a critical role in Fn pathogenicity and interactions with host cells, including cancer cells^36^. Both ATCC strains expressed key virulence genes, including *aim1, fadA, fomA*, and *radD*. Of the eight isolates tested, all carried the *fadA* gene, highlighting its importance in *F. nucleatum* virulence^17^. However, none of the clinical isolates carried *cmpA, fap2*, or *radD*, despite their established roles as virulence factors. Since all primers used in this study were designed based on the genome of strain 23726, it is possible that nucleotide differences between the gene homologs from these isolates were sufficiently different resulting in primers not able to hybridize to the DNA strand. Currently, of all the comparative genomic studies published to date on *F. nucleatum* isolates, only the fadA gene was discussed and was mentioned to be conserved among all *F. nucleatum* isolates while none of the other genes we attempted to amplify here were analyzed. Therefore, it is possible that all of the clinical isolates do carry homologs of these genes. Whole genome sequencing of these isolates are currently underway and we will update the results afterwards. This finding provides the first evidence that many well-studied *F. nucleatum* virulence factors might be absent in some isolates. Particularly notable is the absence of *fap2*, whose product facilitates binding to tumor-expressed Gal-GalNAc. This discrepancy raises questions about alternative virulence mechanisms employed by these isolates and suggests that *F. nucleatum* strains may rely on diverse genetic strategies to adapt to different host environments. Whole genome sequencing of these isolates is currently underway to elucidate the genetic basis of these differences.

Regarding cancer-related effects, the OSCC isolate Co25544 exhibited migration-promoting effects on CRC cells comparable to those of the wild-type strain. However, other clinical isolates demonstrated significantly weaker effects on cell migration. This suggests that Co25544 may possess unique pathogenic factors absent in other isolates. The increased migratory ability of CRC cells following Co25544 infection may also indicate increase ability to promote epithelial-mesenchymal transition (EMT). This is particularly intriguing given that Co25544 lacks the *fap2* gene, a known mediator of CRC progression. These findings imply that *F. nucleatum* isolates may harbor alternative factors that promote tumor progression, warranting further investigation.

Our study demonstrates for the first time that *F. nucleatum* isolates display substantial diversity in cell length, biofilm formation, and the presence of well-studied virulence genes. Importantly, we found that most clinical isolates lacked the ability to promote CRC cell migration, suggesting that this capability may require factors beyond FadA. These findings highlight the need to identify alternative virulence pathways and genetic adaptations in *F. nucleatum*. The absence of *fap2* in clinical isolates, coupled with the robust CRC migration-promoting effects of Co25544, suggests that novel virulence mechanisms may be at play. Future studies should explore the role of unexplored genes and proteins in modulating host-pathogen interactions. Comparative genomic analysis of these clinical isolates will be instrumental in identifying novel virulence factors and understanding strain-specific pathogenicity. Additionally, these findings have implications for diagnostics and therapeutic targeting. The variability in virulence gene profiles underscores the potential for tailored antimicrobial strategies based on strain-specific characteristics. By uncovering new virulence pathways, we may also identify biomarkers for more accurate risk assessment and treatment of *F. nucleatum*-associated diseases. Whole genome sequencing and comparative genomics are currently underway to further elucidate these findings. Future studies will focus on characterizing additional human clinical isolates with the goal of identifying novel virulence factors and expanding our understanding of *F. nucleatum*-host interactions.

## Acknowledgement

We are thankful for Dr. Jenn-Ren Hsiao from the Department of Otolaryngology at National Cheng Kung University Hospital, Taiwan and Dr. Jeffrey S. Chang from the National Institute of Cancer Research at National Health Research Institutes, Taiwan for providing saliva samples. This work was supported by OSU-CHS Startup funding to I-H-H. The funding agency played no role in study design, data collection or analysis, manuscript preparation, or decision to publish.

## Author Contributions

S. L., W-H. H., N-H. W., C. H., C. V., K. S. performed the experiments. S.L., prepared the manuscript. J.-R.H., J. S C., and J.-Y. C. provided saliva samples. J-W. C. contributed to experimental design, co-supervision, and review of this manuscript. I-H.H., contributed to project conceptualization, experimental design, project supervision, preparation and editing of the final manuscript. All authors have read and approved the final version of this manuscript.

## References

1. Fan, Z., et al. *Fusobacterium nucleatum* and its associated systemic diseases: epidemiologic studies and possible mechanisms. J. Oral Microbiol. 15, 2145729 (2023).

2. Parhi, L. et al. Breast cancer colonization by *Fusobacterium nucleatum* accelerates tumor growth and metastatic progression. Nat. Commun. 11, 3259 (2020).

3. Alon-Maimon, T., Mandelboim, O. & Bachrach, G. *Fusobacterium nucleatum* and cancer. Periodontol. 2000 89, 166–180 (2022).

4. Potts, T. V., Holdeman, L. V. & Slots, J. Basic Biological Sciences Relationships Among the Oral Fusobacteria Assessed by DNA-DNA Hybridization. J. Dent. Res. 62, 702–705 (1983).

5. Dzink, J. L., Sheenan, M. T. & Socransky, S. S. Proposal of Three Subspecies of *Fusobacterium nucleatum* Knorr 1922: *Fusobacterium nucleatum* subsp. nucleatum subsp. nov., comb. nov.; Fusobacterium nucleatum subsp. polymorphum subsp. nov., nom. rev., comb. nov.; and Fusobacterium nucleatum subsp. vincentii subsp. nov., nom. rev., comb. nov. Int. J. Syst. Bacteriol. 40, 74–78 (1990).

6. Gharbia, S. E. & Shah, H. N. Heterogeneity within *Fusobacterium nucleatum*, proposal of four subspecies. Lett. Appl. Microbiol. 10, 105–108 (1990).

7. Ye, X., et al. *Fusobacterium nucleatum* Subspecies *Animalis* Influences Proinflammatory Cytokine Expression and Monocyte Activation in Human Colorectal Tumors. Cancer Prev. Res. (Phila. Pa.) 10, 398–409 (2017).

8. Alluri, L. S. C. et al. Presence of Specific Periodontal Pathogens in Prostate Gland Diagnosed With Chronic Inflammation and Adenocarcinoma. Cureus (2021) doi:10.7759/cureus.17742.

9. Socransky, S. S., Haffajee, A. D., Cugini, M. A., Smith, C. & Kent, R. L. Microbial complexes in subgingival plaque. J. Clin. Periodontol. 25, 134–144 (1998).

10. Chukkapalli, S. S. et al. Polymicrobial Oral Infection with Four Periodontal Bacteria Orchestrates a Distinct Inflammatory Response and Atherosclerosis in ApoEnull Mice. PLOS ONE 10, e0143291 (2015).

11. Šaferis, V. Relationship of Clinical and Microbiological Variables in Patients with Type 1 Diabetes Mellitus and Periodontitis. Med. Sci. Monit. 20, 1871–1877 (2014).

12. Han, Y. W., et al. *Fusobacterium nucleatum* Induces Premature and Term Stillbirths in Pregnant Mice: Implication of Oral Bacteria in Preterm Birth. Infect. Immun. 72, 2272–2279 (2004).

13. Strauss, J. et al. Invasive potential of gut mucosa-derived *Fusobacterium nucleatum* positively correlates with IBD status of the host: *Inflamm*. Bowel Dis. 17, 1971–1978 (2011).

14. Williams, M. D., Kerber, C. A. & Tergin, H. F. Unusual Presentation of Lemierre’s Syndrome Due to *Fusobacterium nucleatum*. J. Clin. Microbiol. 41, 3445–3448 (2003).

15. Sparks Stein, P., et al. Serum antibodies to periodontal pathogens are a risk factor for Alzheimer’s disease. Alzheimers Dement. 8, 196–203 (2012).

16. Al-hebshi, N. N. et al. Inflammatory bacteriome featuring *Fusobacterium nucleatum* and Pseudomonas aeruginosa identified in association with oral squamous cell carcinoma. Sci. Rep. 7, 1834 (2017).

17. Rubinstein, M. R., et al. *Fusobacterium nucleatum* Promotes Colorectal Carcinogenesis by Modulating E-Cadherin/β-Catenin Signaling via its FadA Adhesin. Cell Host Microbe 14, 195–206 (2013).

18. Da, J., Wang, X., Li, L. & Xu, Y. *Fusobacterium nucleatum* Promotes Cisplatin-Resistance and Migration of Oral Squamous Carcinoma Cells by Up-Regulating Wnt5a-Mediated NFATc3 Expression. Tohoku J. Exp. Med. 253, 249–259 (2021).

19. Gholizadeh, P., Eslami, H. & Kafil, H. S. Carcinogenesis mechanisms of *Fusobacterium nucleatum*. Biomed. Pharmacother. 89, 918–925 (2017).

20. Sun, J., et al. *F. nucleatum* facilitates oral squamous cell carcinoma progression via GLUT1-driven lactate production. eBioMedicine 88, 104444 (2023).

21. Li, Y. et al. Intracellular *Fusobacterium nucleatum* infection attenuates antitumor immunity in esophageal squamous cell carcinoma. Nat. Commun. 14, 5788 (2023).

22. Shang, F.-M. & Liu, H.-L. *Fusobacterium nucleatum* and colorectal cancer: A review. World J. Gastrointest. Oncol. 10, 71–81 (2018).

23. Flanagan, L., et al. *Fusobacterium nucleatum* associates with stages of colorectal neoplasia development, colorectal cancer and disease outcome. Eur. J. Clin. Microbiol. Infect. Dis. 33, 1381–1390 (2014).

24. Mima, K., et al. *Fusobacterium nucleatum* in colorectal carcinoma tissue and patient prognosis. Gut 65, 1973– 1980 (2016).

25. Gur, C. et al. Binding of the Fap2 Protein of *Fusobacterium nucleatum* to Human Inhibitory Receptor TIGIT Protects Tumors from Immune Cell Attack. Immunity 42, 344–355 (2015).

26. Xu, M. et al. FadA from *Fusobacterium nucleatum* Utilizes both Secreted and Nonsecreted Forms for Functional Oligomerization for Attachment and Invasion of Host Cells. J. Biol. Chem. 282, 25000–25009 (2007).

27. Han, Y. W. et al. Identification and Characterization of a Novel Adhesin Unique to Oral Fusobacteria. J. Bacteriol. 187, 5330–5340 (2005).

28. Coppenhagen-Glazer, S. et al. Fap2 of *Fusobacterium nucleatum* Is a Galactose-Inhibitable Adhesin Involved in Coaggregation, Cell Adhesion, and Preterm Birth. Infect. Immun. 83, 1104–1113 (2015).

29. Yu, T., et al. *Fusobacterium nucleatum* Promotes Chemoresistance to Colorectal Cancer by Modulating Autophagy. Cell 170, 548–563.e16 (2017).

30. Ramos, A. & Hemann, M. T. Drugs, Bugs, and Cancer: *Fusobacterium nucleatum* Promotes Chemoresistance in Colorectal Cancer. Cell 170, 411–413 (2017).

31. Tomkovich, S. et al. Locoregional Effects of Microbiota in a Preclinical Model of Colon Carcinogenesis. Cancer Res. 77, 2620–2632 (2017).

32. Kapatral, V. et al. Genome Sequence and Analysis of the Oral Bacterium *Fusobacterium nucleatum* Strain ATCC 25586. J. Bacteriol. 184, 2005–2018 (2002).

33. Krieger, M. et al. Stratification of *Fusobacterium nucleatum* by local health status in the oral cavity defines its subspecies disease association. Cell Host Microbe 32, 479–488.e4 (2024).

34. Weiser, J. N. The battle with the host over microbial size. Curr. Opin. Microbiol. 16, 59–62 (2013).

35. Yang, D. C., Blair, K. M. & Salama, N. R. Staying in Shape: the Impact of Cell Shape on Bacterial Survival in Diverse Environments. Microbiol. Mol. Biol. Rev. 80, 187–203 (2016).

36. Chen, Y. et al. More Than Just a Periodontal Pathogen –the Research Progress on *Fusobacterium nucleatum*. Front. Cell. Infect. Microbiol. 12, 815318 (2022).

37. Lima, B. P., Shi, W. & Lux, R. Identification and characterization of a novel *Fusobacterium nucleatum* adhesin involved in physical interaction and biofilm formation with *Streptococcus gordonii*. MicrobiologyOpen 6, e00444 (2017).

